# Functional Compensation Between Hematopoietic Stem Cells *In Vivo*

**DOI:** 10.1101/236398

**Authors:** Lisa Nguyen, Zheng Wang, Elizabeth Chu, Jiya Eerdeng, Adnan Y. Chowdhury, Rong Lu

**Affiliations:** Eli and Edythe Broad Center for Regenerative Medicine and Stem Cell Research, Department of Stem Cell Biology and Regenerative Medicine, Keck School of Medicine, University of Southern California, Los Angeles, CA 90089, USA

## Abstract

In most organ systems, regeneration is a coordinated effort that involves many stem cells, but little is known about whether and how individual stem cells compensate for the functional deficiencies of other stem cells. Functional compensation between stem cells is critically important during disease progression and treatment. Here, we show how individual hematopoietic stem cells (HSCs) in a mouse heterogeneously compensate for the deficiencies of other HSCs’ during lymphopoiesis by increasing their clonal expansion at specific differentiation stages. This compensation rescues the overall blood supply and influences blood cell types outside of the deficient lineages in distinct patterns. We have identified the molecular regulators and signaling pathways in HSCs that are involved in this process. Our data demonstrate how stem cells interact with each other to constitute a coordinated network that is robust enough to withstand minor functional disruptions. Exploiting the innate compensation capacity of stem cell networks may improve the prognosis and treatment of many diseases.

## Introduction

Hematopoietic stem cells (HSCs) are widely scattered throughout the body in dispersed bone marrow niches^1,2^. Yet they must work together to maintain a common pool of peripheral blood cells. We have recently shown that HSCs can adapt their differentiation programs to the presence of other HSCs at various transplantation doses to ensure overall balanced blood production^3^. A similar coordination mechanism may allow healthy HSCs to rescue the functional deficiencies of impaired HSCs.

It is critically important to understand how the functional deficiencies of a subset of HSCs impact the overall HSC network. Many hematopoietic diseases arise from either an abnormal abundance (i.e., myeloproliferative disorder, thrombocytosis, leukocytosis, and erythrocytosis) or an abnormal deficiency (i.e., myelodysplastic syndrome, neutropenia, agranulocytosis, and thrombocytopenia) of certain blood cell types^4–6^. The initial stages of these diseases may involve a latent period when a patient’s healthy HSCs can sufficiently compensate for the deficiencies of diseased cells and ameliorate disease symptoms. Moreover, the primary treatment for many of these diseases, bone marrow transplantation^7^, also requires donor HSCs to adapt their differentiation programs to the disease environment. HSC compensation, especially in the lymphoid lineages, may play a role in aging as it is common for lymphopoietic deficiencies to develop in the elderly^8,9^. Very few studies have attempted to understand the innate compensation capacity of stem cell networks. In this study, we offer new insights on HSC compensation mechanisms at the cellular and molecular levels.

While a single HSC is capable of regenerating the entire blood and immune system under extreme circumstances^10–12^, limiting dilution assays of HSC transplantation show that the number of donor HSCs quantitatively determines the fraction of blood cells that they produced^13,14^. This suggests a simple coordination model, where individual HSCs play equal roles in contributing to blood regeneration after transplantation. Recent work from our group and others have shown that HSC differentiation is heterogeneous at the clonal level^10,15–19^. For example, individual HSC clones supply differential amounts of blood cells in mice and in human patients ^17,18,20,21^. Moreover, recent studies of native hematopoiesis suggest that different blood cell types have distinct clonal origins^22^. These new findings raise the question of how the diverse differentiation programs of individual HSCs are coordinated in the aftermath of functional disruptions. In this study, we use a co-transplantation experimental model and high throughput genetic barcode tracking technology to simultaneously quantify the *in vivo* differentiation of many individual HSCs. We show how normal HSCs compensate for genetically mutated HSCs that are deficient in supplying one or multiple types of lymphocytes (Fig 1).

**Figure 1:**
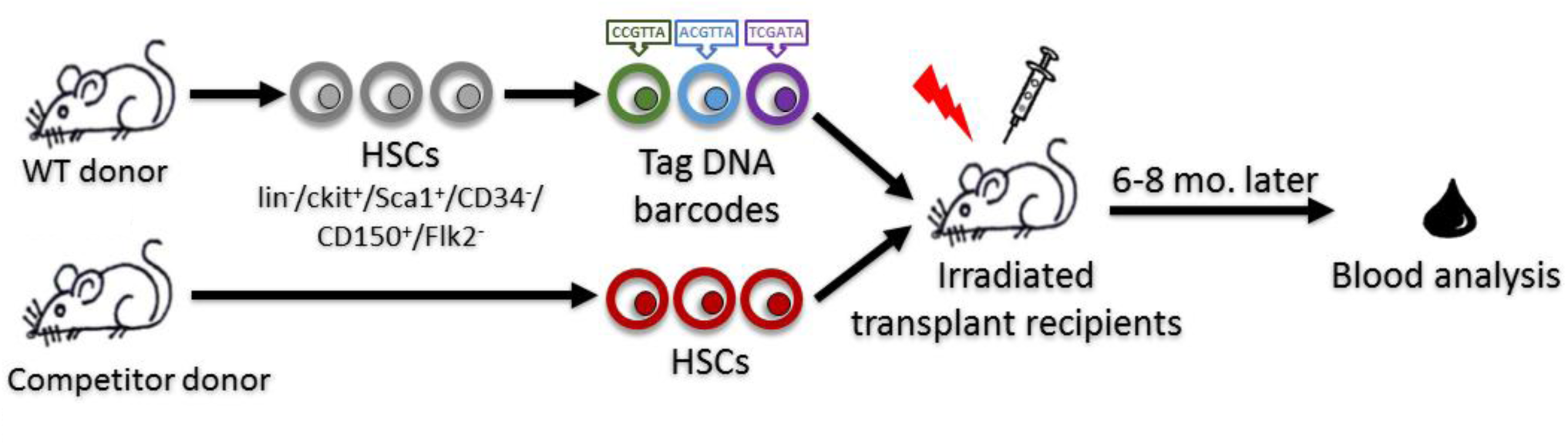
A co-transplantation mouse model to investigate stem cell compensation. Schematic depicting co-transplantation of barcoded wildtype (WT, CD45.2) HSCs with either WT (CD45.1), B cell deficient (uMT ^-/-^, CD45.1), or B and T cell deficient (Rag2^-/-^γc^-/-^ or NSG, CD45.1) HSCs into irradiated recipient mice (CD45.1/CD45.2). Peripheral blood is harvested from recipient mice at 6-8 months post transplantation for population and clonal level analyses.

## Results

To determine if HSCs functionally compensate for each other within an organism, we co-transplanted wild type (WT) HSCs with lineage-deficient HSCs into WT recipient mice (Fig 1). Lineage-deficient HSCs were isolated from uMT^-/-^ mice that are unable to produce B cells, and from NSG and Rag2^-/-^γc^-/-^ mice that are unable to produce both B and T cells (Fig 2B, 2E, Appendix Figure S1B, 1E, EV Fig 1). We examined the peripheral blood of the recipient mice 6-8 months post transplantation when blood production had returned to a steady state (Fig 1)^14,23^. We found that WT HSCs compensated for the deficiencies of their partner HSCs in distinct patterns. In the uMT^-/-^ co-transplantation group, WT HSCs significantly oversupplied B cells (Fig 2A, Appendix Figure S1A). This suggests that the presence of lineage-deficient HSCs changed the differentiation of WT HSCs within a common host. Moreover, we found that the presence of WT HSCs also changed the differentiation of lineage-deficient HSCs. For example, uMT^-/-^ mice have normal levels of T cells (EV Fig 1), but uMT^-/-^ HSCs produced fewer T cells in the co-transplantation setting (Fig 2B, Appendix Figure S1B). The reduction in B and T cell production by uMT^-/-^ HSCs was compensated by an oversupply from WT HSCs such that the overall B and T cell levels remained normal (Fig 2C, Appendix Figure S1C). In the Rag2^-/-^γc^-/-^ and NSG co-transplantation groups, WT HSCs significantly oversupplied B cells, CD4 T and CD8 T cells (Fig 2D, Appendix Figure S1D) to maintain overall B and T cell production (Fig 2F, Appendix Figure S 1F). Thus, our data suggest that WT HSCs increased their differentiation specifically in the deficient lineages to maintain the overall lymphocyte supply.

**Figure 2:**
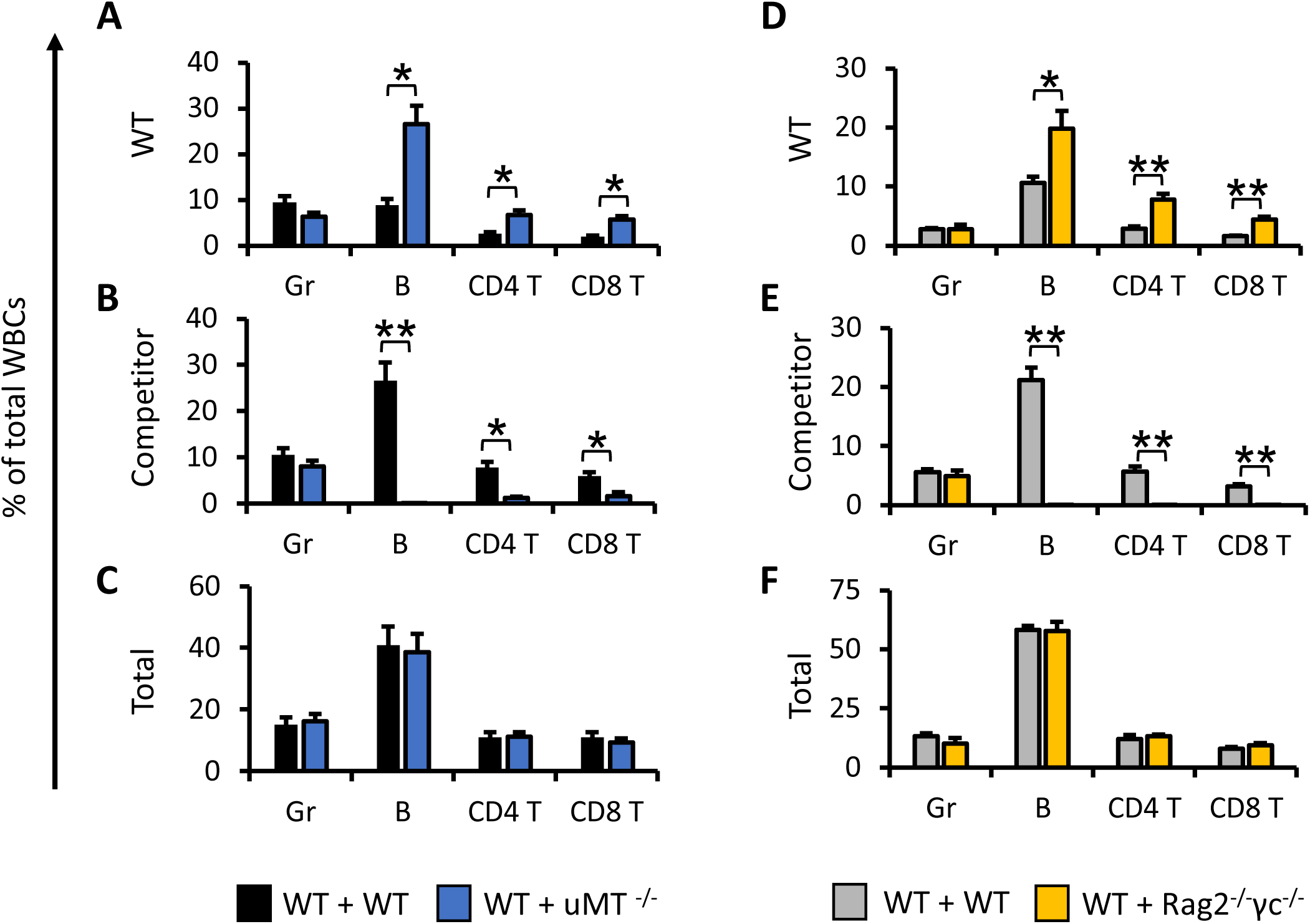
HSCs compensate for the lineage deficiencies of co-transplanted HSCs in blood production. A-F The production of granulocytes (Gr), B cells, CD4 T cells, and CD8 T cells in the peripheral blood by WT donor HSCs, competitor donor HSCs (uMT ^-/-^ and Rag2^-/-^γc^-/-^HSCs), and the total cell production are shown as percentages of the total number of white blood cells (WBCs). Data information: Data were collected 6 months post transplantation and presented as mean ± SEM. *p < 0.05 (Student’s t-test). n = 8 mice for each group.

To determine if lymphopoietic compensation originates from an increase in the number of differentiating HSCs or from an elevated expansion of their differentiation, we used a genetic barcoding technique that we have developed to track WT donor HSCs at the clonal level (Fig 1)^17^. We found that the total numbers of WT HSC clones that supply the myeloid and lymphoid lineages were similar between the control and co-transplantation groups (Fig 3, Appendix Figure S2). To compare HSC clonal expansion, we classified HSC clones by the quantity of their blood cell production. We found that significantly more HSC clones produced high amounts of B cells in the uMT^-/-^ co-transplantation group, and high amounts of B and T cells in the NSG and Rag2^-/-^γc^-/-^ co-transplantation group (Fig 4B, 4F-H, Appendix Figure S3B, 3F-H). The increase in clonal expansion was not observed in the granulocyte population (Fig 4A, 4E, Appendix Figure S3A, 3E). This suggests that more HSCs highly expand in the specific lineages where compensation is needed. In addition, we also found that the levels of their expansion increased during compensation. The most abundant clones supplied significantly larger amounts of lymphocytes in the corresponding deficient lineages (Fig 4I-J, Appendix Figure S3I-J). Taken together, our data suggest that lineage deficiency is primarily compensated for by highly expanded HSC clones and not by an increase in the total number of differentiating HSC clones.

**Figure 3:**
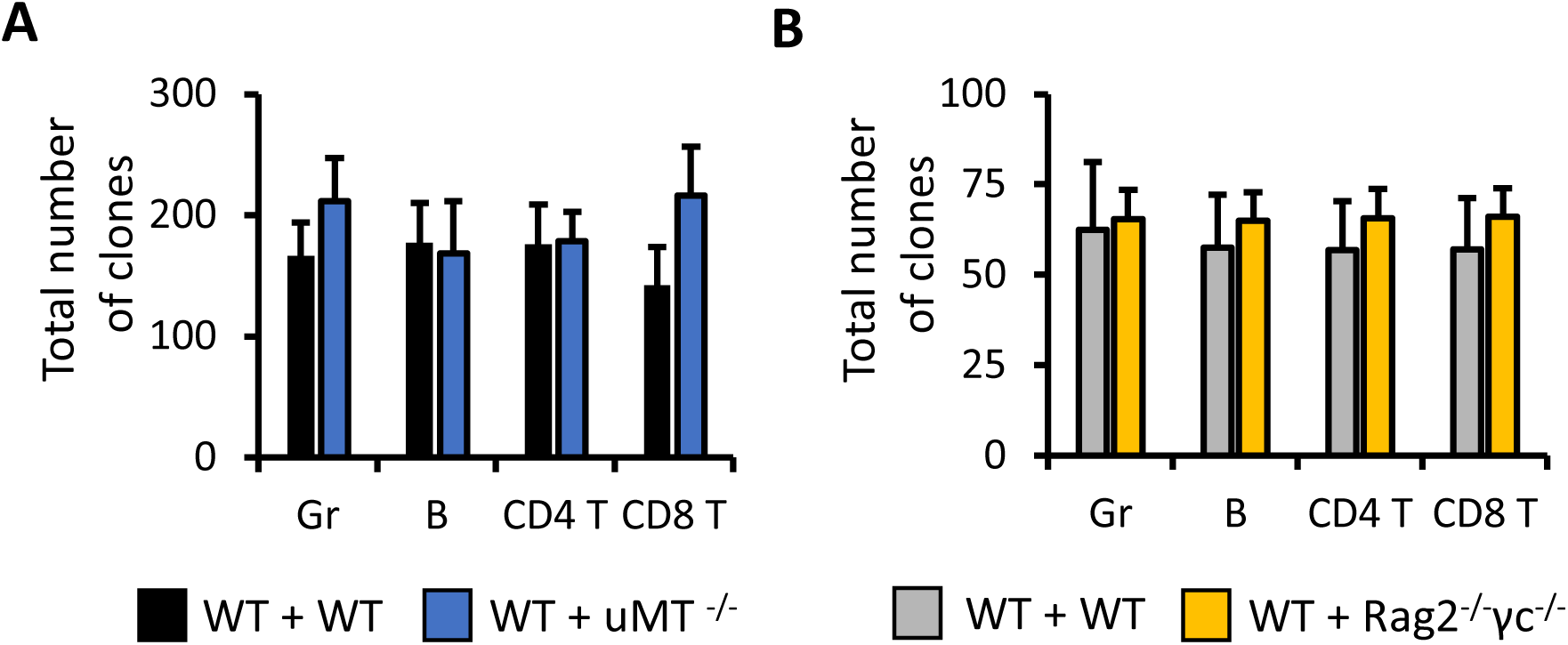
The number of differentiating HSC clones does not change in the presence of lineage deficient HSCs. A-B Total numbers of barcoded clones that give rise to granulocytes (Gr), B cells, CD4 T cells, and CD8 T cells. Data information: Data were collected 6 months post transplantation and presented as mean ± SEM. n = 8 mice for each group.

**Figure 4:**
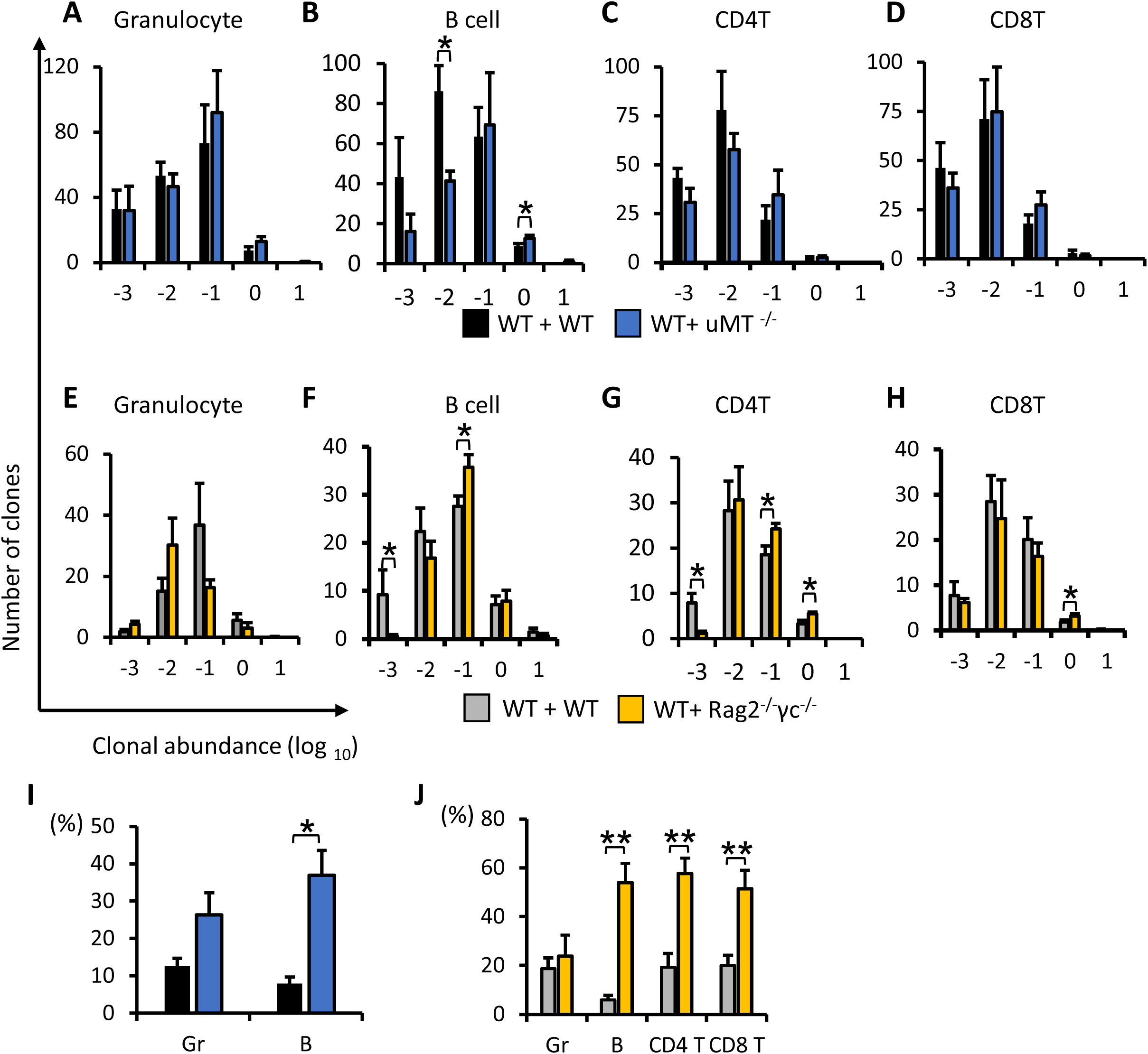
Highly expanded HSC clones increase their differentiation in response to the lineage deficiencies of other HSCs. A-H Number of HSC clones that produce different amounts of granulocytes (Gr), B cells, CD4 T cells, and CD8 T cells. I-J Production of Gr, B, CD4 T, and CD8 T by the top 5 most abundant clones in each lineage. Data information: Data were collected 6 months post transplantation and presented as mean ± SEM. *P < 0.05 (Student’s t-test). n = 8 mice for each group.

HSC differentiation is a step-wise process, during which HSCs expand dramatically in number. To identify the differentiation stage at which compensation takes place, we analyzed the number of intermediate progenitors (Fig 5), particularly those derived from WT donor HSCs (Appendix Figure S4). In both co-transplantation groups, we found a significant increase in the number of common lymphoid progenitors (CLPs) (Fig 5, Appendix Figure S4). As CLPs supply B and T cells, an increase in CLPs to compensate for B cell deficiency will increase T cell production as well. This explains the T-cell compensation that we observed in the uMT^-/-^ co-transplantation group (Fig 2A-C, Appendix Figure S1A-C). In addition to an increase of the lymphoid progenitors, the uMT^-/-^ co-transplantation group also exhibited a reduction in myeloid progenitors such as common myeloid progenitors (CMPs) and granulocyte macrophage progenitors (GMPs) (Fig 5A, Appendix Figure S4A). Surprisingly, there are significantly fewer HSCs in both co-transplantation groups, indicating that HSC differentiation compensation may be associated with compromised self-renewal. The decrease in HSC abundance may have resulted in reduced levels of multipotent progenitors (MPPs) (Fig 5, Appendix Figure S4).

**Figure 5:**
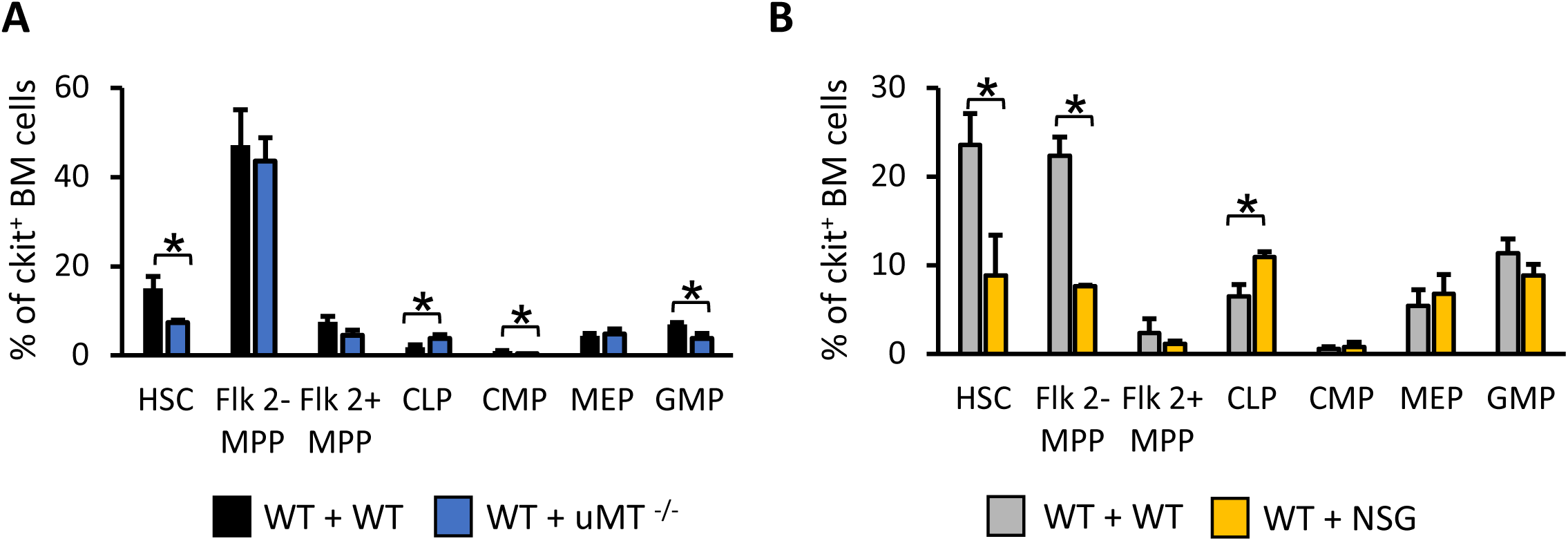
Compensation for lineage deficiency occurs at the progenitor level. A-B Abundance of progenitors as a percentage of ckit+ bone marrow (BM) cells (n = 6 mice for each group). Shown are hematopoietic stem cells (HSCs), Flk2- and Flk2+ multipotent progenitors (MPP), common lymphoid progenitors (CLP), common myeloid progenitors (CMP), megakaryocyte-erythroid progenitors (MEP), and granulocyte-monocyte progenitors (GMP) Data information: Data were collected at 7 months post transplantation for uMT-/- group and 8 months post transplantation for NSG group, and presented as mean ± SEM. *p < 0.05 (Student’s t-test). n = 6 mice for each group.

To determine whether the lymphoid compensation is associated with any molecular level changes in HSCs, we compared the gene expression profiles of WT HSCs from the uMT^-/-^ and NSG co-transplantation groups with WT HSCs from the control groups 7-8 months after transplantation. We found that compensating HSCs significantly changed their gene expression profile (Fig 6, Appendix Figure S5, EV Fig 2-5). In particular, 96 significantly changed genes are shared between the uMT^-/-^ and NSG groups (Fig 6A). Genes that change similarly between the uMT^-/-^ and NSG co-transplantation groups have been found to be expressed in hematopoietic populations and play roles in regulating lymphopoiesis (Fig 6B). For example, PAX5 is a master regulator of B cell differentiation. It represses B-lineage-inappropriate genes and activates B-lymphoid-specific genes^24^. PTPRO is induced by Wnt/β-catenin signaling in a lymphoid enhancer factor dependent manner^25^. IGF1 has been shown to regulate primary B lymphopoiesis^26^. Up-regulation of these genes may be instrumental in priming HSCs to compensate for lymphoid lineages.

**Figure 6:**
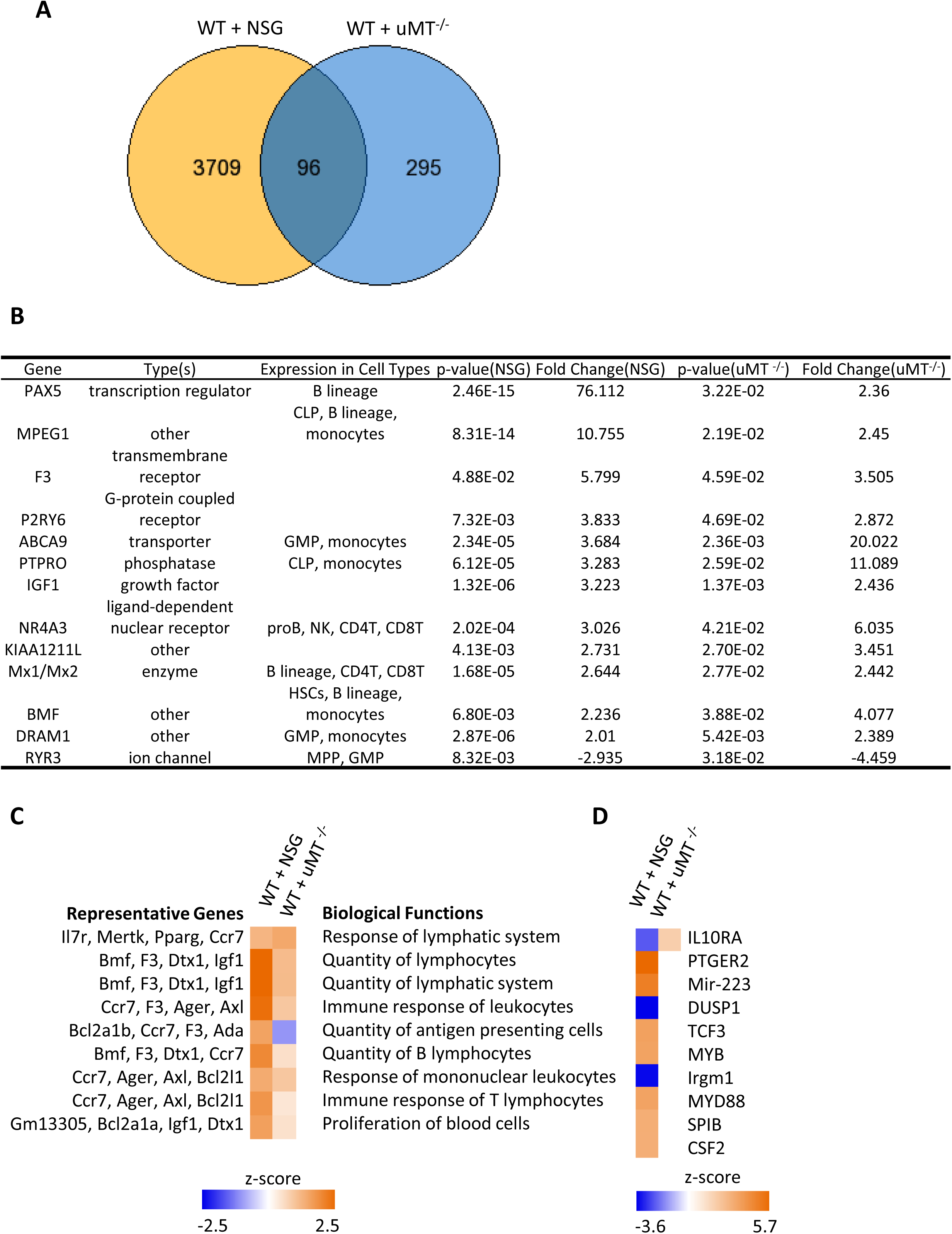
Differential gene expression of HSCs during their compensation for lineage deficiency. A Number of genes that are differentially expressed in WT HSCs co-transplanted with lineage deficient HSCs compared to control co-transplanted with WT HSCs. Gene lists are generated by Partek Flow. Thresholds are defined as p < 0.05 and fold change < 2 or > -2. B Top genes whose expression changed similarly in the NSG and uMT^-/-^ co-transplantation groups. This gene list is generated by Ingenuity Pathway Analysis (IPA). C Top functions that are activated or inactivated, generated by IPA. Shown are the functions with Fisher’s Exact Test p-value < 0.05. D Predicted upstream regulators that are activated or inactivated, generated by IPA. Data information: Data were collected at 7 months post transplantation for uMT^-/-^ group and 8 months post transplantation for NSG group. n = 3 mice for each group, except for the NSG control group where n = 2 mice are analyzed. The activation z scores correspond to a statistical measure of the match between expected relationship direction and observed gene expression.

To identify biological functions of the differentially expressed genes, we performed Ingenuity Pathway Analysis (IPA) (Fig 6C-D). Among the genes that are differentially expressed in both uMT^-/-^ and NSG groups, we found significant activation of functions related to response, quantity, and proliferation of lymphocytes (Fig 6C, Appendix Figure S5). The degree of activation is generally higher in the NSG group. We applied the same analysis to each co-transplantation group separately and found that the highest scored diseases, biological functions, and networks are those that are relevant to lymphoid compensation (EV Fig 3-4). In addition, we identified interleukin 10 receptor alpha subunit (IL10RA) as an upstream regulator whose expression appears to decrease with NSG and increase with uMT^-/-^ co-transplantation (Fig 6D). The prediction is based on the comparison of IL10RA downstream target genes and the differentially expressed genes (EV Fig 5). IL10RA encodes a lymphoid cytokine receptor that has been shown to mediate the immunosuppressive signal of interleukin 10 and inhibit the synthesis of proinflammatory cytokines^27^.

## Discussion

In summary, we have presented an experimental model suitable for studying the functional compensation between individual HSCs *in vivo.* By transplanting different types of stem cells into a single recipient, our mouse model allows for investigating the interactions between stem cells. Similar approaches can be applied to study other tissue and organ systems and to other disease models. Our data demonstrate how the HSC network operates *in vivo* as a robust coordinated system. The compensation capacities of individual HSCs enable the HSC network to tolerate partial loss of function. Some HSCs quantitatively expand their differentiation to compensate for deficient HSCs and specifically overproduce undersupplied cell types. This suggests that individual HSCs heterogeneously respond to the differentiation deficiencies of other HSCs. This heterogeneous compensation may be essential for maintaining robustness in blood regeneration and suggests that stem cell coordination is a complex process. Furthermore, we have identified cellular and molecular mechanisms underlying the functional compensation between HSCs. In particular, compensation for lymphoid deficiencies is observed at the CLP level and influences the differentiation in other downstream lymphoid lineages. While cellular level compensation manifests itself at the progenitor level, molecular changes have been identified at the stem cell level. We have discovered molecular regulators and pathways in HSCs that are associated with the compensation. Future studies can manipulate these regulatory molecules to improve the efficacy of bone marrow transplantation and to develop new therapeutic strategies exploiting endogenous HSC compensation capacity. New prognosis approaches may be developed by monitoring the compensation activities of endogenous HSCs. A better understanding of stem cell interactions can help improve the treatment of many degenerative and age-related diseases.

## Methods

### Mice

WT donor mice used in the co-transplantation experiments were C57BL/6J (CD45.2). The lineage deficient donor mice were B6.129S2(B6)-*Ighm^tm1Cgn^*/J (uMT^-/-^, CD45.1), NOD-*scid* IL2Rgamma^null^ (NSG) and C;129S4-*Rag2^tm1.1Flv^ Il2rg^tm1.1Flv^*/J (Rag2^-/-^γc^-/-^, CD45.1). The recipient mice were offsprings of C57BL/6J and B6.SJL-Ptprca Pepcb/BoyJ (F1, CD45.1/ CD45.2). All donor and recipient mice were 8-12 weeks old at the time of transplantation. Irradiation was performed on all recipient mice before transplantation at 950 cGy. We examined 5-8 mice for each experimental group, and performed biological replicates shown in supplemental figures. Mice were bred and maintained at the Research Animal Facility of the University of Southern California. Animal procedures were in compliance with the International Animal Care and Use Committee.

### Cell Isolation and Transplantation

HSCs (lineage (CD3, CD4, CD8, B220, Gr1, Mac1, Ter119)-/ckit+/Sca1+/Flk2-/CD34-/CD150+) were obtained from the crushed bones of donor mice and isolated using FACS sorting with the FACS-Aria II (BD Biosciences, San Jose, CA) after enrichment using CD117 microbeads (AutoMACS, Miltenyi Biotec, Auburn, CA). HSCs were infected for 15 hours with lentivirus carrying barcodes and then transplanted via retro-orbital injection. HSC clonal labeling was performed as described previously^17^. Whole bone marrow cells (helper cells) were flushed from the femurs of F1 mice (*CD45.1/ CD45.2*) mice and added to HSCs right before transplantation. 250,000 helper cells were transplanted per mouse. For both single deficient and double deficient experiments, we co-transplanted barcoded WT HSCs with uMT^-/-^ HSCs (1:3 and 1:2 ratios) and with Rag2^-/-^γc^-/-^ and NSG HSCs (1:2 ratio). For the control group, we co-transplanted HSCs from WT (CD45.1) and WT (CD45.2) donor mice at the same ratio as the experimental group.

### Blood Sample Collection and FACS Analysis

Blood samples were collected into PBS containing 10 mM EDTA via a small transverse cut in the tail vein. To eliminate red blood cells, 2% dextran was added, and the remaining blood cells were treated with ammonium-chloride-potassium lysis buffer on ice for 5 minutes to remove residual red blood cells. After a 30-minute antibody incubation at 4° C, samples were suspended in PBS with 2% FBS and 4,6-Diamidino-2-phenylindole to distinguish dead cells. Cells were sorted using the FACS-Aria I and II cell sorters and separated into granulocytes, B cells, CD4 T cells, and CD8 T cells. Antibodies were obtained from eBioscience and BioLegend as described previously^17^. Donor cells were sorted based on the CD45 marker. The following cell surface markers were used to harvest hematopoietic populations:

Granulocytes: CD4-/CD8-/B220-/CD19-/Mac1+/Gr1+/side scatter^high^;
B cells: CD4-/CD8-/Gr1-/Mac1-/B220+/CD19+;
CD4 T cells: B220-/CD19-/Mac1-/Gr1-/TCRαβ+/CD4+/CD8-;
CD8 T cells: B220-/CD19-/Mac1-/Gr1-/TCRαβ+/CD4-/CD8+;
HSC (hematopoietic stem cells): lineage (CD3, CD4, CD8, B220, Gr1, Mac1, Ter119)-/ /ckit+/Sca1+/Flk2-/CD34-/CD150+

Flow cytometry data were analyzed using FlowJo software version 9.6.1 (Tree Start, Ashland, OR) and Diva software.

### DNA Barcode Extraction and Sequencing

Genomic DNA (gDNA) was extracted from sorted hematopoietic cells and amplified using Phusion PCR mastermix (Thermo Scientific, Waltham, MA). The PCR reactions were halted once they had progressed halfway through the exponential phase. PCR product was purified and analyzed using high-throughput sequencing at the USC Epigenome Center. Sequencing data were analyzed as described^3,17^. We combined sequencing data with FACS data to generate the clonal abundance for each HSC clone as the percentage of white blood cells.

Clonal abundance =100%* (Donor % in the blood cell population) * (GFP % among donor cells) * (number of reads for each barcode) / (total reads of all barcodes)

### RNA Sequencing

Total RNA was isolated from HSCs using the Zymo Research (Irvine, CA) Quick-RNA MicroPrep. A separate mouse was used for each triplicate sample. RNA concentration and integrity was determined by BioAnalyzer 2100(Agilent Technologies) using Agilent RNA 6000 Pico Kit. And libraries were constructed according to the manufacturer’s instructions using SMARTer stranded total RNAseq kit or SMART-Seq Ultra Low input RNA Kit (TaKaRa). Each sample was sequenced to produce 40-60 million 75bp single-end reads at the University of Southern California Epigenome Center and Children’s Hospital Los Angeles Genomics Core. Data is analyzed by Partek^®^ Flow^®^ software, version 6.0 Copyright©^27^. After adaptor trimming and quality control to treat the raw reads, the clean reads were aligned to mm10 genome and transcriptome reference using STAR 2.5, default parameters. Gene counts module in Partek pipeline was used to quantification gene expression, excluding those whose maximum raw read counts were less than or equal to 10 counts. Differentially expressed genes are identified by the Partek E/M method with fold changes below -2 and greater than 2, and with a P value less than 0.05.

The networks and functional analyses were generated using Ingenuity Pathway Analysis (IPA)^29^. The biological functional analysis is based on expected causal effects between genes, which are derived from the literature knowledge base of IPA. The analysis examines genes in the dataset that are known to affect biological functions. Functions with fewer than 10 differentially expressed genes are excluded from the list. The threshold for all biological functions bar graphs is –log (p-value) = 1.3, which is calculated using the Fisher’s exact test right-tailed. Predictions of activation or inactivation of functions and pathways were based on the z-score algorithm, which takes into account the causal relationships between genes and biological functions and networks determined by the direction of effect.

## Acknowledgements

We thank USC Libraries Bioinformatics Service for assisting with data analysis. The bioinformatics software and computing resources used in the analysis are funded by the USC Office of Research and the Norris Medical Library. We thank A. Nogalska and Q. Liu for helpful discussions and C. Lytal for manuscript edits. We also thank Y. Chen, M. Li, and T Trecek for help with RNA sequencing analysis; A. Nogalska for laboratory management; L. Barsky, J. Boyd, and B. Masinsin for FACS core management; and C. Nicolet for RNA processing and high-throughput sequencing. This work is supported by NIH R00-HL113104, R01HL135292, R01HL138225 and P30CA014089. L.N. is supported by an NIH T32 training grant and F31 HL134359. E.C. is supported by the Rose Hills Foundation Science and Engineering Fellowship and the USC Provost’s Undergraduate Research Fellowship. A.C. is supported by the California Institute for Regenerative Medicine Training Grant and the Hearst Fellowship Award.

## Author Contributions

L.N. and R.L. designed and performed the experiments. E.C. and J.E. wrote custom Python codes for data analysis. Z.W. and A.C. assisted with RNA experiments. L.N. and R.L. wrote the manuscript. All authors edited the manuscript.

---

**Appendix Figure S1:**
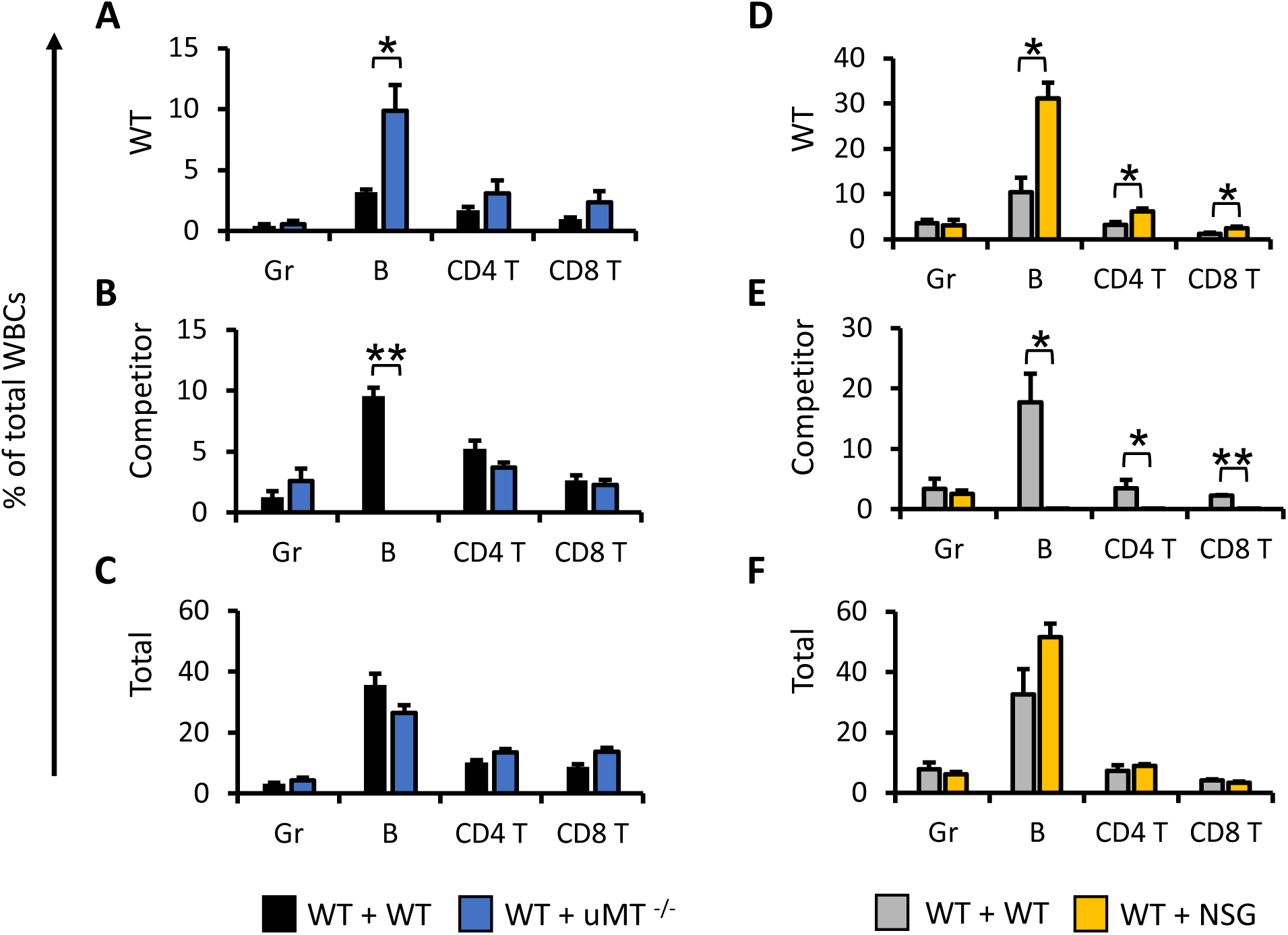
HSCs compensate for the lineage deficiencies of co-transplanted HSCs in blood production (replicate experiment of Figure 2). A-F The production of granulocytes (Gr), B cells, CD4 T cells, and CD8 T cells in the peripheral blood by WT donor HSCs, competitor donor HSCs (uMT ^-/-^ and NSG HSCs), and the total cell production are shown as percentages of the total number of white blood cells (WBCs). Data information: Data were collected at 7 months post transplantation for uMT^-/-^ group and 8 months post transplantation for NSG group, and presented as mean ± SEM. *p < 0.05 (Student’s t-test). n = 6 mice for each group.

**Appendix Figure S2:**
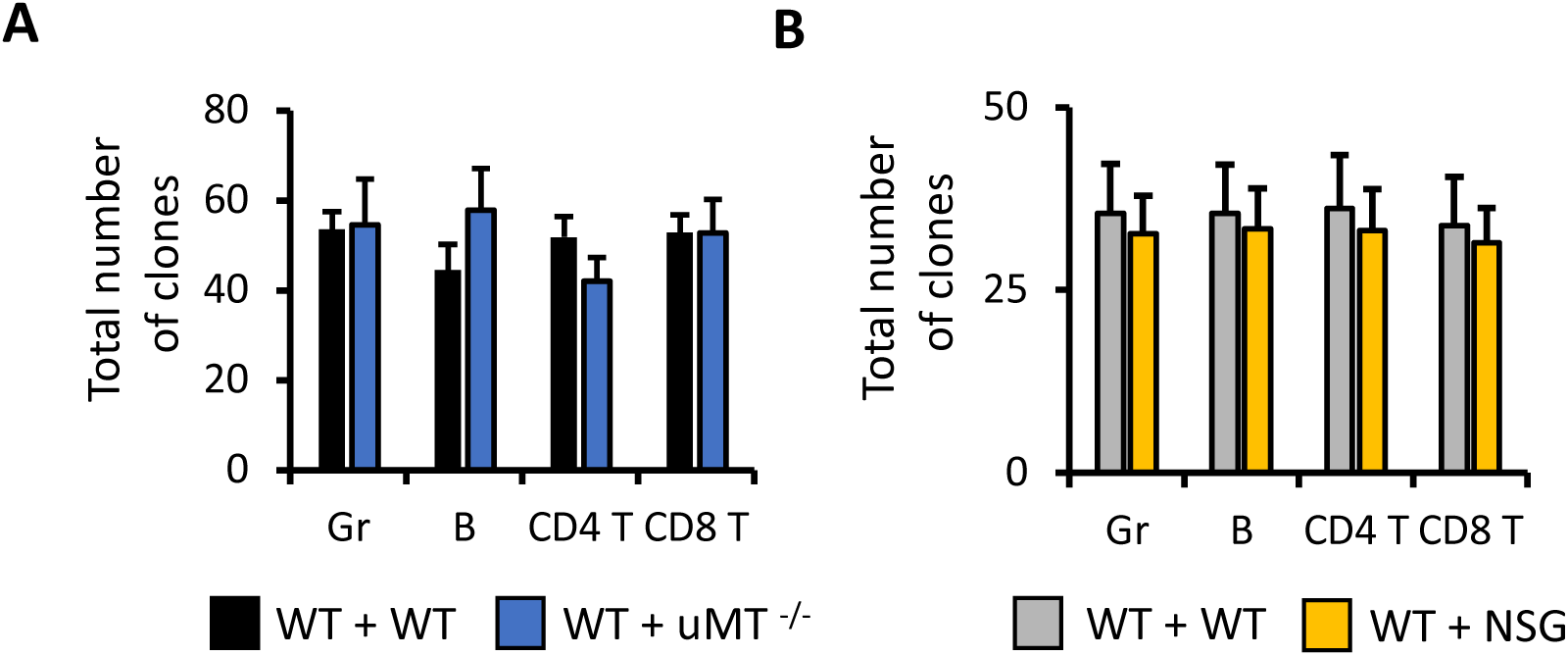
The number of differentiating HSC clones does not change in the presence of lineage deficient HSCs (replicate experiment of Figure 3). A-B Total numbers of barcoded HSC clones that give rise to granulocytes (Gr), B cells, CD4 T cells, and CD8 T cells. Data information: Data were collected at 7 months post transplantation for uMT^-/-^ group and 8 months post transplantation for NSG group, presented as mean ± SEM. n = 6 mice for each group.

**Appendix Figure S3:**
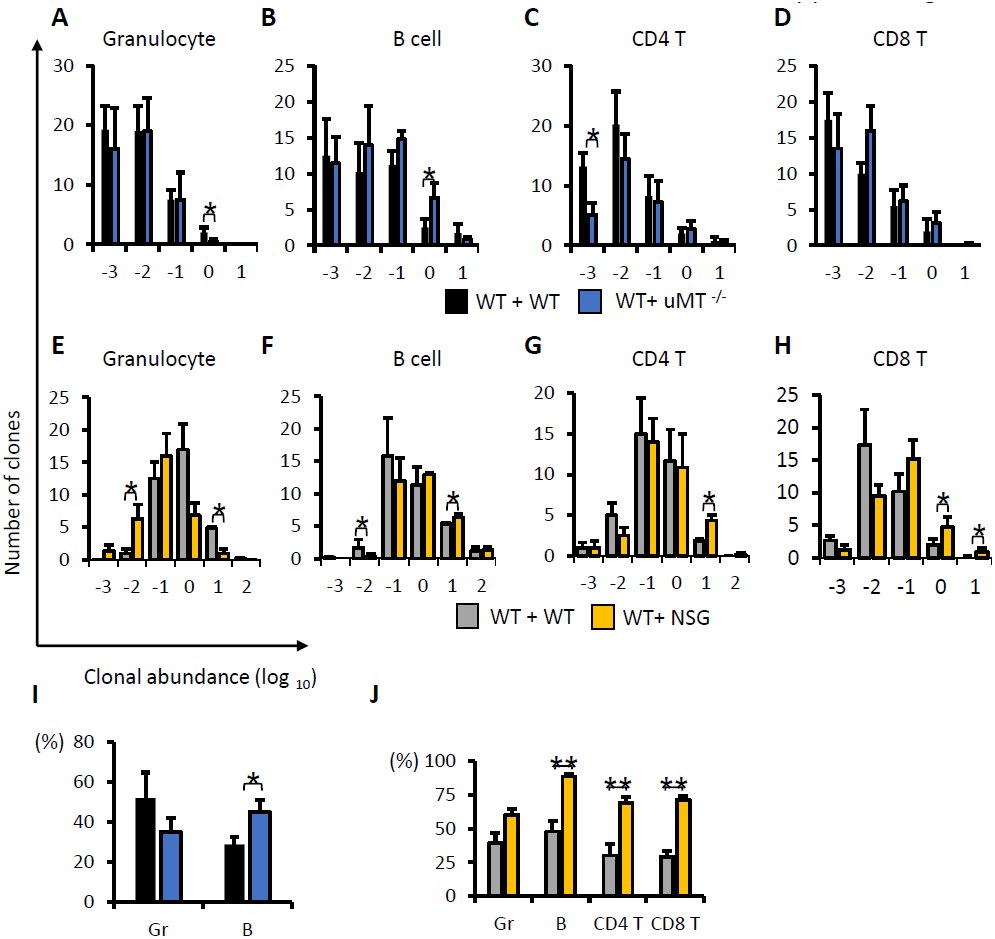
Highly expanded HSC clones increase their differentiation in response to the lineage deficiencies of other HSCs (replicate experiment for Figure 4). A-H Number of HSC clones that produce different amounts of granulocytes (Gr), B cells, CD4 T cells, and CD8 T cells. I-J Production of Gr, B, CD4 T, and CD8 T by the top 5 most abundant clones in each lineage. Data information: Data were collected at 7 months post transplantation for uMT^-/-^ group and 8 months post transplantation for NSG group, and presented as mean ± SEM. *p < 0.05 (Student’s t-test). n = 6 mice for each group.

**Appendix Figure S4:**
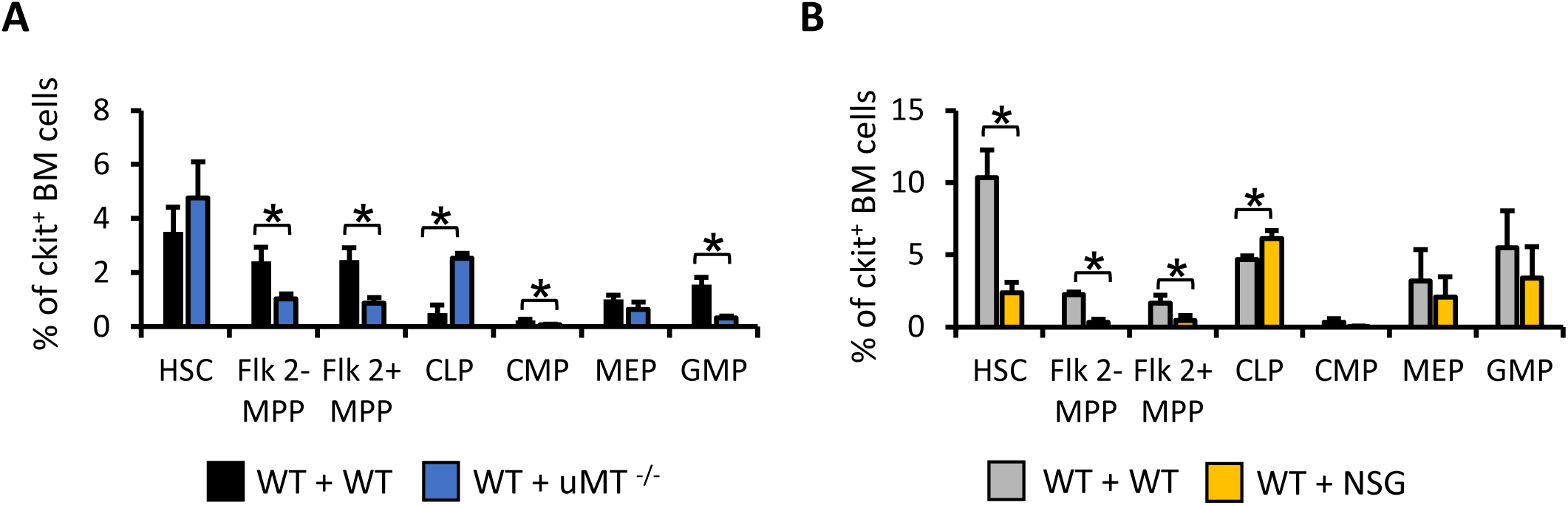
Compensation for lineage deficiency occurs at the progenitor level. A-B Abundance of WT donor HSC derived progenitors as a percentage of ckit+ bone marrow (BM) cells (n = 6 mice for each group). Shown are hematopoietic stem cells (HSCs), Flk2- and Flk2+ multipotent progenitors (MPP), common lymphoid progenitors (CLP), common myeloid progenitors (CMP), megakaryocyte-erythroid progenitors (MEP), and granulocyte-monocyte progenitors (GMP) Data information: Data were collected at 7 months post transplantation for uMT^-/-^ group and 8 months post transplantation for NSG group, and presented as mean ± SEM. *p < 0.05 (Student’s t-test). n = 6 mice for each group.

**Appendix Figure S5:**
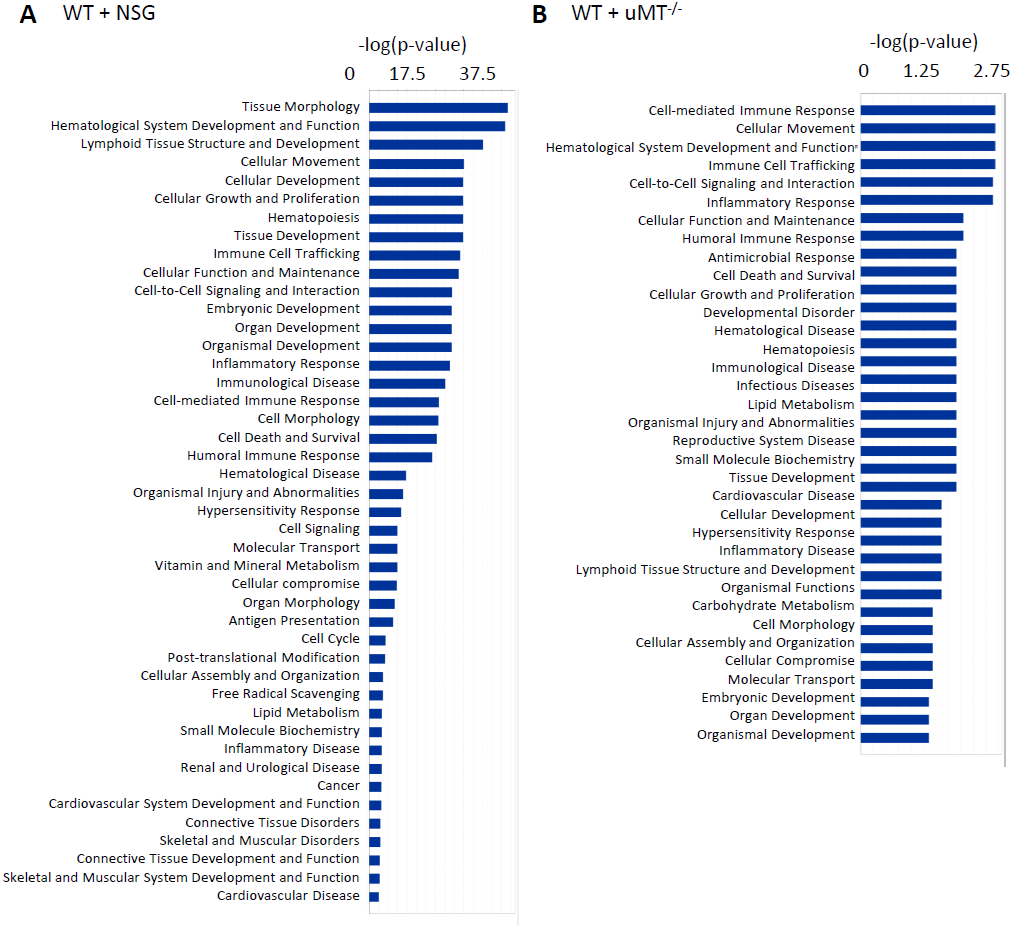
All diseases and functions that are activated in the presence of deficient HSCs (Fisher’s Exact Test p-value < 0.05).

**Extended View Figure 1:**
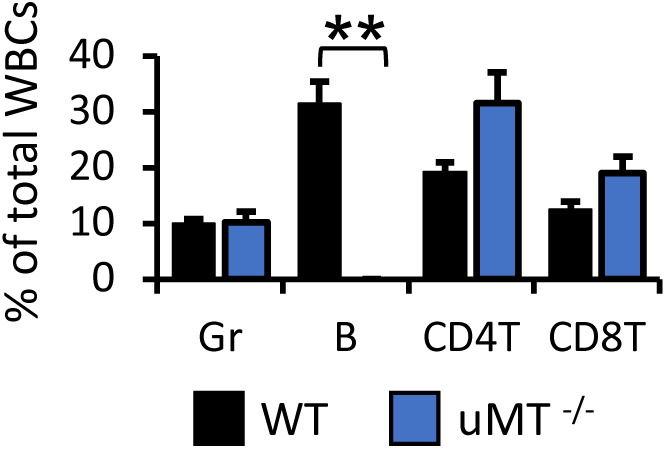
Lineage-deficient donor HSCs do not supply all blood cell types. uMT ^-/-^ mice do not produce B cells, but produce normal levels of granulocytes (Gr), B cells, CD4 T cells, and CD8 T cells. **p < 0.005 (Student’s t-test).

**Extended View Figure 2:**
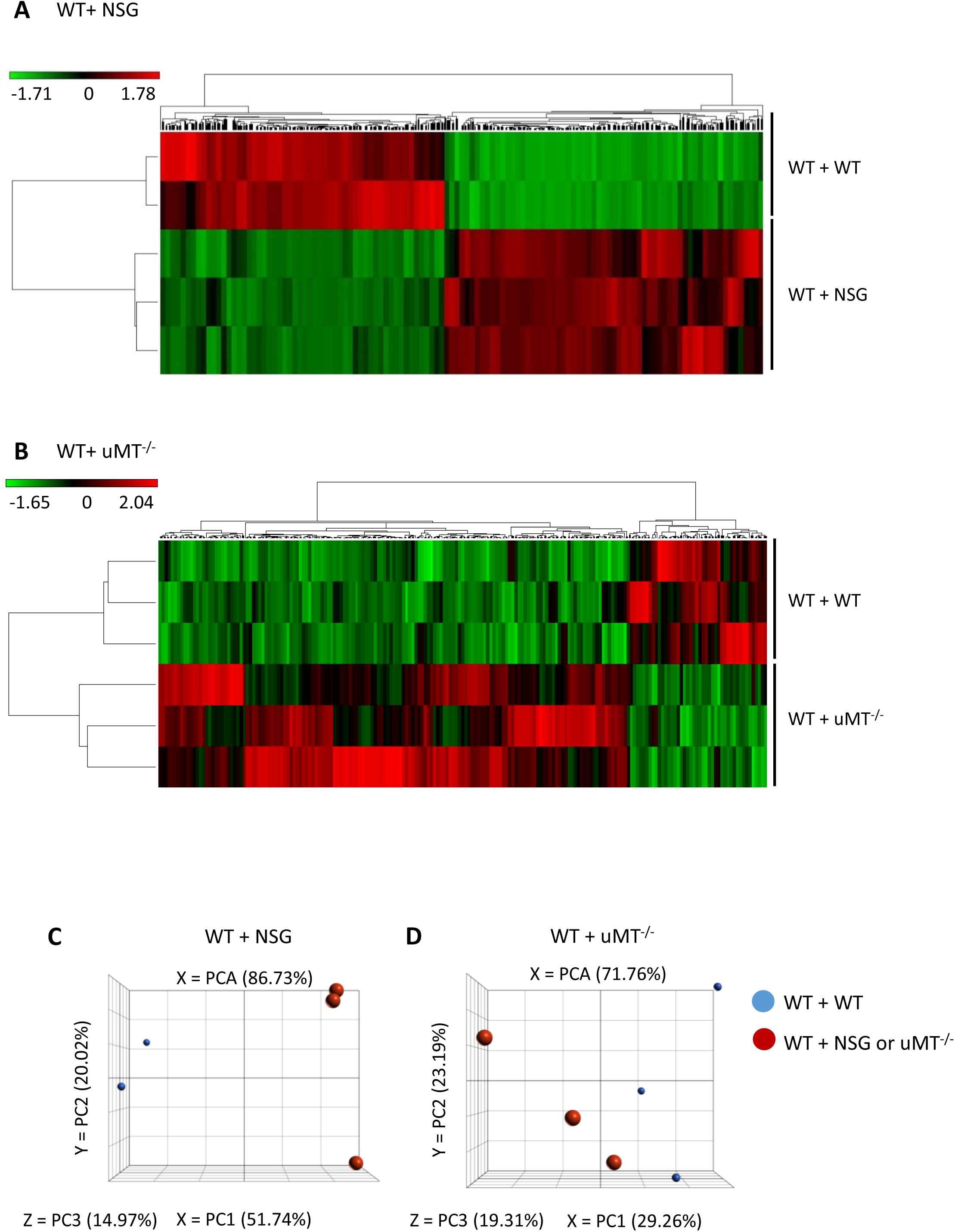
Biological replicate samples are clustered together based on gene expression. A-B Unsupervised clustering of individual samples using average linkage for cluster and Pearson correlation for point distance metrics. Data information: Data were collected 7 months post transplantation. n = 3 mice for each group. C-D Principal component analysis of individual samples in control and treatment groups. Data information: Data were collected at 7 months post transplantation for uMT^-/-^ group and 8 months post transplantation for NSG group. n = 3 mice for each group, except for the NSG control group where n = 2 mice are analyzed.

**Extended View Figure 3:**
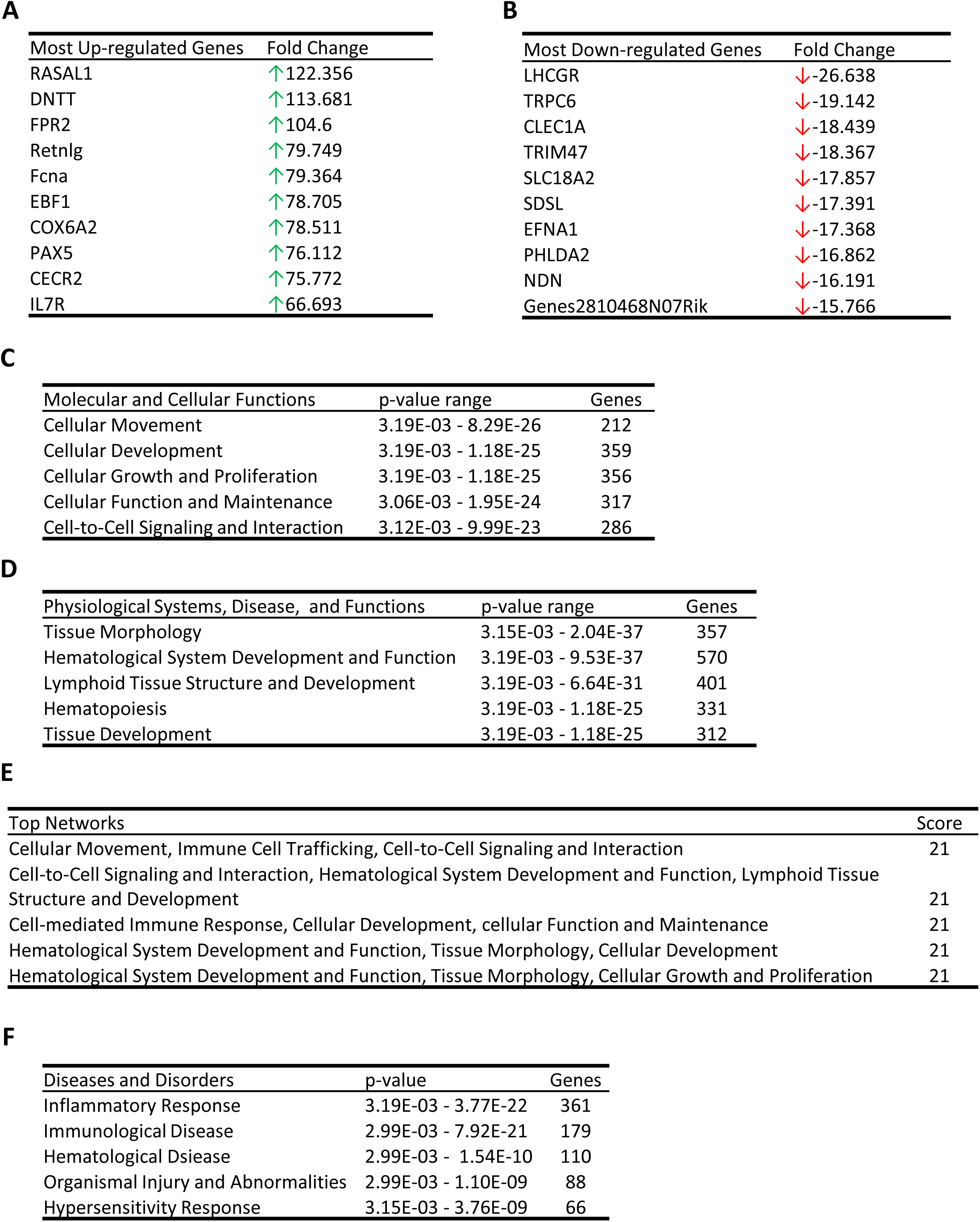
Gene expression signature of WT HSCs co-transplanted with NSG HSCs. A-B Top genes that are differentially expressed. C Top molecular and cellular functions that are activated. D Top physiological systems, diseases and functions that are significantly affected. E Top networks that are significantly affected. The scores take into account the number of focus genes and the size of the network to approximate the relevance of the network to the original list of genes. F Top diseases and disorders that are significantly affected. Data information: Ingenuity Pathway Analysis (IPA) was used to analyze differentially expressed genes. The activation z scores correspond to a statistical measure of the match between expected relationship direction and observed gene expression and are displayed using a dark blue to orange gradient.

**Extended View Figure 4:**
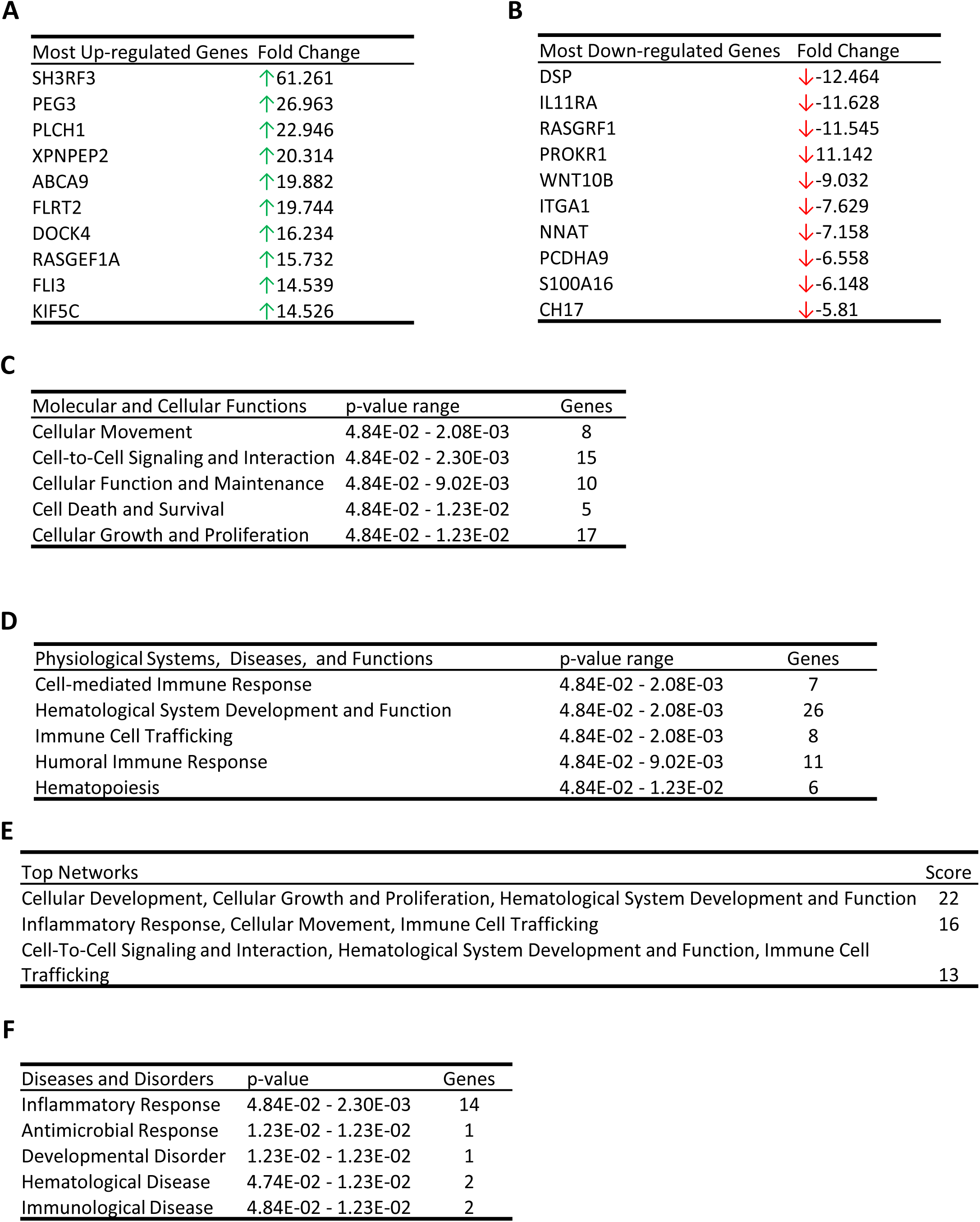
Gene expression signature of WT HSCs co-transplanted with uMT^-/-^ HSCs. A-B Top genes that are differentially expressed. C Top molecular and cellular functions that are activated. D Top physiological systems, diseases and functions that are significantly affected. E Top networks that are significantly affected. The scores take into account the number of focus genes and the size of the network to approximate the relevance of the network to the original list of genes. F Top diseases and disorders that are significantly affected. Data information: IPA was used to analyze differentially expressed genes. The activation z scores correspond to a statistical measure of the match between expected relationship direction and observed gene expression and are displayed using a dark blue to orange gradient.

**Extended View Figure 5:**
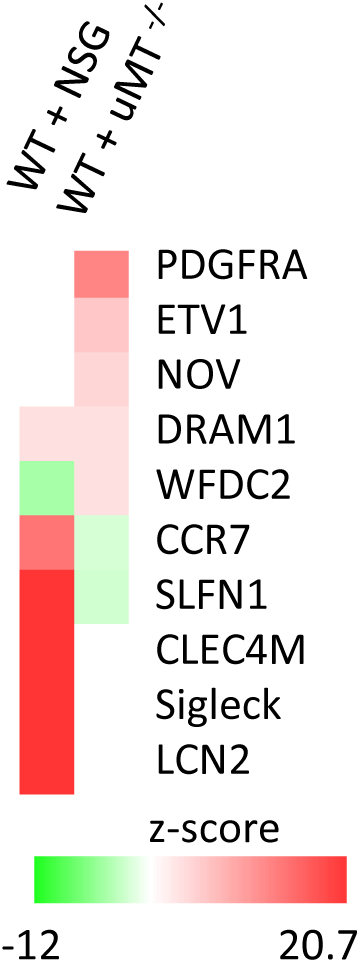
Genes downstream of IL10RA that are differentially expressed in the co-transplantation groups.

